# Ammonia-oxidizing archaea and bacteria differentially contribute to ammonia oxidation in soil under precipitation gradients and land legacy

**DOI:** 10.1101/2023.11.08.566028

**Authors:** Soumyadev Sarkar, Anna Kazarina, Paige M. Hansen, Kaitlyn Ward, Christopher Hargreaves, Nicholas Reese, Qinghong Ran, Willow Kessler, Ligia F.T. de Souza, Terry D. Loecke, Marcos V. M. Sarto, Charles W. Rice, Lydia H. Zeglin, Benjamin A. Sikes, Sonny T.M. Lee

## Abstract

**Background:** Global change has accelerated the nitrogen cycle. Soil nitrogen stock degradation by microbes leads to the release of various gases, including nitrous oxide (N_2_O), a potent greenhouse gas. Ammonia-oxidizing archaea (AOA) and ammonia-oxidizing bacteria (AOB) participate in the soil nitrogen cycle, producing N_2_O. There are outstanding questions regarding the impact of environmental processes such as precipitation and land use legacy on AOA and AOB structurally, compositionally, and functionally. To answer these questions, we analyzed field soil cores and soil monoliths under varying precipitation profiles and land legacies.

**Results:** We resolved 28 AOA and AOB metagenome assembled genomes (MAGs) and found that they were significantly higher in drier environments and differentially abundant in different land use legacies. We further dissected AOA and AOB functional potentials to understand their contribution to nitrogen transformation capabilities. We identified the involvement of stress response genes, differential metabolic functional potentials, and subtle population dynamics under different environmental parameters for AOA and AOB. We observed that AOA MAGs lacked a canonical membrane-bound electron transport chain and F-type ATPase but possessed A/A-type ATPase, while AOB MAGs had a complete complex III module and F-type ATPase, suggesting differential survival strategies of AOA and AOB.

**Conclusions:** The outcomes from this study will enable us to comprehend how drought-like environments and land use legacies could impact AOA– and AOB-driven nitrogen transformations in soil.

## Background

Human activities as well as global climate change have significantly altered the global nitrogen cycle within the past century [1]. Nitrogen (N) inputs from industrial sources and crop N fixation exceed inputs from natural N fixation [2]. Similarly, the global N cycle has accelerated significantly as a positive feedback with climate change [3]. Soil N stocks represent the largest organic terrestrial pool of N, and microbial degradation of soil N stocks can lead to substantial releases of greenhouse gases (GHGs), including nitrous oxide (N_2_O) [4]. N_2_O is a potent GHG with a global warming potential 265 times that of CO_2_ [5]. N_2_O is the most important stratospheric ozone-depleting substance [6]. This is particularly alarming, as microbial sources of N_2_O may shift with environmental changes [4]. To reduce N_2_O emissions, we need a thorough understanding of biogeochemical pathways and environmental variables that shape the microbial functions underpinning the N cycle and N_2_O dynamics [4].

Nitrification, the microbial oxidation of ammonia to nitrite (NO_2−_) and nitrate (NO_3−_), is a central step in the N cycle and provides substrates for N removal processes and ammonia oxidation to N_2_ [7,8]. Ammonia oxidation, which is the first step in nitrification, is performed by three groups of ammonia-oxidizing microorganisms (AOOs): ammonia-oxidizing bacteria (AOB), ammonia-oxidizing archaea (AOA), and complete ammonia oxidizers (comammox *Nitrospira*) [9–12]. AOA and AOB not only directly produce N_2_O but also provide substrate for N_2_O production through denitrification [4,13]. AOA and AOB may play a pivotal, underexplored role in fueling denitrification and facilitating terrestrial N_2_O emissions in many soil environments [13]. However, little is known about the environmental conditions that favor each process and thereby N_2_O production and consumption [4].

AOA often dominate AOB in pristine neutral and acidic soils [14–16], while in most fertilized agricultural soils, AOB are deemed the predominant ammonia oxidizers [11,12,16–18]. However, their significance in agricultural systems and how they are affected by global climate change remain poorly understood. To date, most studies on the AOA and AOB use selective genes and assays to understand the relative abundance and to discern the activities of the microbial populations in the environment [19–23]. Although these pioneering studies have provided insights into the potential nitrification processes of AOA and AOB, mechanistic insights into the functional controls on the nitrification activities of either group remain scarce [24].

Ammonium concentration [18,25,26], pH [15,16,22], salinity [27–29], and dissolved concentration of oxygen [29–31] are important factors that control the AOA and AOB community structures and composition. Precipitation is also another component that can influence AOA and AOB communities, with many nitrifiers being found to be drought-resistant and responsive to rewetting [32]. However, studies have only begun to understand the functions of AOA and AOB influenced by changing precipitation profiles. Therefore, there is a clear knowledge gap with a need for more efforts to decipher the functional insights under varying gradients of precipitation and different land legacies.

In this study, we analyzed 18 soil cores in the field and 36 cores in soil monoliths across different land legacy and precipitation gradients to determine how AOA and AOB community structure and composition are regulated across precipitation gradients and land legacy. The use of both the soil cores and monoliths complemented each other – the soil monoliths enhanced the current precipitation from the soil cores, which harbored long legacies of precipitation and land use; that may alter the function of AOOs to show the inertia of legacy effects compared to current precipitation in the field. As such, we hypothesized that AOA would harbor specific survival strategies that contribute to their adaptability in drier environmental conditions compared to AOB populations. We were also interested in deciphering the functional mechanistic insights that were instrumental in driving the contribution of those microbes associated with ammonia oxidation under differential land legacy.

## Methods

### Sample collection, monolith experiment, and processing

Figure 1 shows the experimental design of our study. In June 2018, we used a Giddings probe (Giddings Machine Company, Windsor, CO, USA) to obtain 60 cm deep (5 cm diameter) soil core samples (n=3) from each of 3 sites in Western (WKS) and Eastern (EKS) Kansas. Sites in western Kansas experience an average of 533 mm/year precipitation, and sites in eastern Kansas have an average rainfall of 1045 mm/year [33]. At both WKS and EKS, we obtained soil cores from 3 different land-use legacies: WKS (native/N: 38.84°N, 99.30°W, agricultural/Ag: 38.84°N, 99.31°W, and postagricultural/PAg: 38.84°N, 99.32°W) and EKS (native/N: 38.18°N, 95.27°W, agricultural/Ag: 38.54°N, 95.25°W, and postagricultural/PAg: 38.18°N, 95.27°W). Within 3 plots in each precipitation region/land use legacy combination, we collected 1 soil core from 3 subplots. Soil cores from subplots were pooled to create 1 composite sample per plot. We subsampled 25 g of soil for shotgun sequencing, and the samples were stored at –80°C until downstream processing.

**Figure 1.**
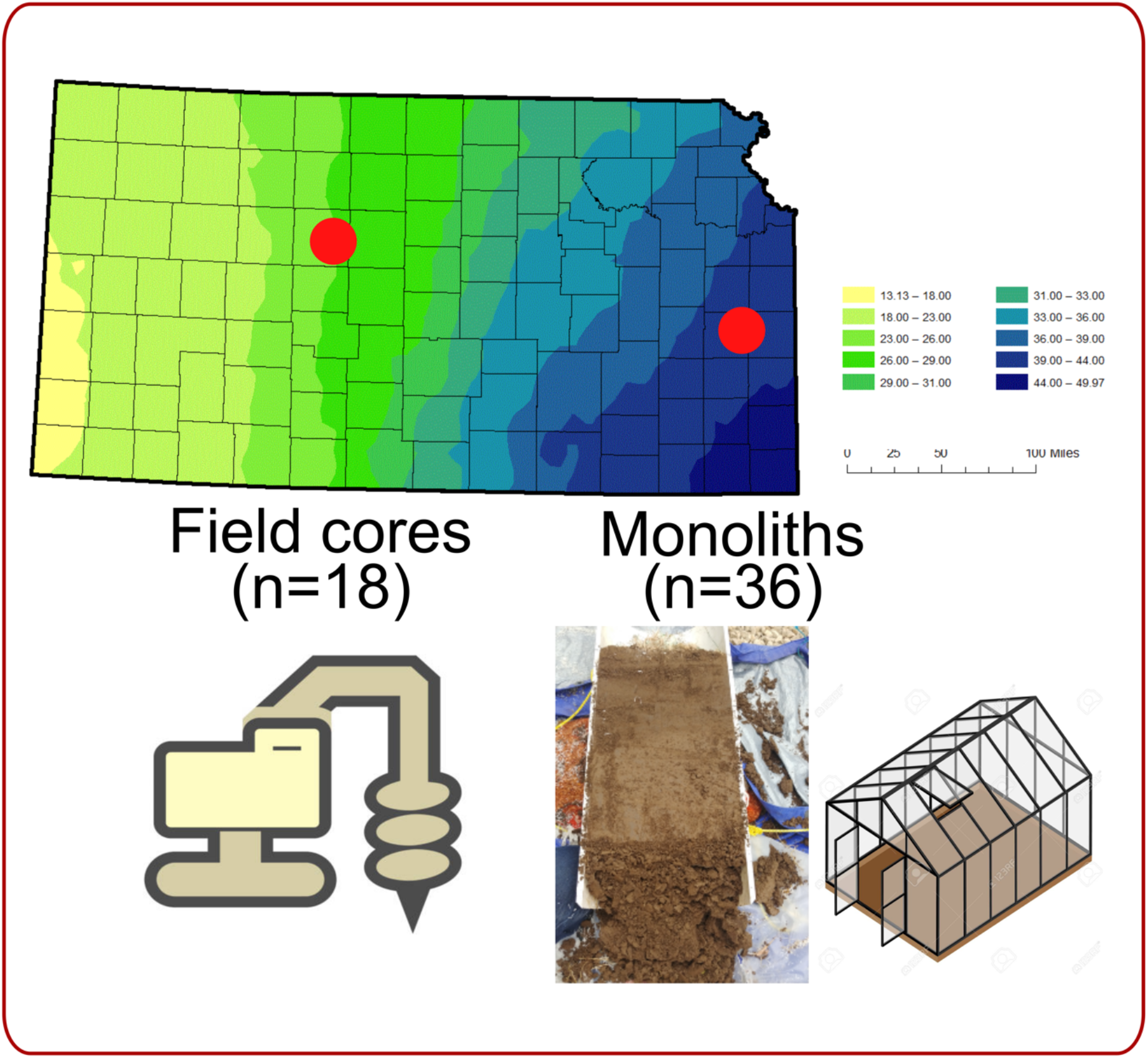
The experimental design of the study highlights the field soil cores and monoliths that were obtained from Western and Eastern Kansas under three different land legacies: agricultural, native, and postagricultural.

From August to October 2018, we also used custom hydraulic machinery [34] to collect a total of 36 (30 cm x 60 cm) intact soil cores from the same sites described above (i.e., 6 soil cores/monoliths per precipitation/land use legacy combination). These monoliths were stored and dried under ambient conditions in a greenhouse at the University of Kansas Field Station until the experiment was set up in April 2019. Each monolith was rewetted and then randomly assigned into ‘wet’ and ‘dry’ watering treatments. For both the EKS and WKS monoliths, watering treatment amounts were determined by averaging 30 years of rainfall data for Hays, KS, then raising that number by 50% to account for increased transpiration demand due to high temperatures in the greenhouse (1000 mm/yr). The wet treatment received twice this amount (2000 mm/yr). Of these total irrigation quantities, 450 mm/yr of each treatment was applied during three 150 mm intense rainfall events per year to mimic summer storm events common across Kansas. After applying these treatments for 6 months, all monoliths were destructively harvested in November 2019. We retained the top 5 cm (25 g of soil) from the monolith for this study and stored it at –80°C until downstream processing.

For the field samples, we homogenized the soil samples before the extractions using the DNeasy PowerSoil Pro Kit following the manufacturer’s protocol (Qiagen, Germantown, MD, USA). E.Z.N.A. A Soil DNA Kit (Omega Biotek, Inc., Norcross, GA, United States) was used to extract microbial DNA from the monoliths following the manufacturer’s protocol with necessary modifications. We measured 0.160 g of monolith soil samples and combined the techniques of bead-beating (20 rev/s, 2 mins) and vortexing (3 mins) to elute the purified DNA to a volume of 100 µl. We assessed the DNA quality, purity, and integrity of the field samples and monoliths by a Nanodrop Spectrophotometer (Thermo Fisher Scientific, MA, USA) and Qubit 4 Fluorometer (Thermo Fisher Scientific, MA, USA). The Illumina NovaSeq 6000 platform (Illumina, San Diego, CA, USA) was used to generate the shotgun metagenomic sequencing for the field samples and the monoliths. We used an S1 flow cell to undertake a 150-paired sequencing strategy using Nextera DNA Flex library preparation.

### Bioinformatic workflow

We organized the dataset into 4 “metagenomic sets” – 2 from the field cores and 2 from the monolith based on the geographic locations (WKS, EKS). We employed coassembly strategies, where we coassembled reads from each metagenomic set. We automated our metagenomics bioinformatics workflows using the program ‘anvi-run-workflow’ in anvi’o v7.1 [35]. The workflows use Snakemake [36] and implement numerous tasks, including short-read quality filtering, assembly, gene calling, functional annotation, hidden Markov model search, metagenomic read recruitment, and binning.

We used the program ‘iu-filer-quality-minoche’ to process the short metagenomic reads and remove low-quality reads according to the criteria outlined previously [37]. We used MEGAHIT v1.2.9 to assemble quality-filtered short reads into longer contiguous sequences (contigs) [38]. We then used the following strategies to process the assemblies: we used (1) ‘anvi-gen-contigs-database’ on contigs to compute k-mer frequencies and identify open reading frames (ORFs) using Prodigal v2.6.3 [39]; (2) ‘anvi-run-hmms’ to identify sets of bacterial and archaeal single-copy core genes using HMMER v.3.2.1 [40]; (3) ‘anvi-run-ncbi-cogs’ to annotate ORFs from NCBI’s Clusters of Orthologous Groups (COGs); and (4) ‘anvi-run-kegg-kofams’ to annotate ORFs from KOfam HMM databases of KEGG orthologs.

We mapped metagenomic short reads to contigs using Bowtie2 v2.3.5 [41] and converted them to BAM files using samtools v1.9 [42]. We profiled the BAM files using ‘anvi-profile’ with a minimum contig length of 1000 bp. We used ‘anvi-merge’ to combine all profiles into an anvi’o merged profile for all downstream analyses. We then used ‘anvi-cluster-contigs’ to group contigs into initial bins using CONCOCT v1.1.0 [43] and used ‘anvi-refine’ to manually curate the bins based on tetranucleotide frequency and different coverage across the samples. We marked bins that were more than 70% complete and less than 10% redundant as metagenome-assembled genomes (MAGs). Finally, we used ‘anvi-compute-genome-similarity’ to calculate the average nucleotide identity (ANI) of our genomes using PyANI v0.2.9 [44] and identified nonredundant MAGs. We used the “detection” metric to assess the occurrence of MAGs in a given sample. Detection of the MAGs is the proportion of the nucleotides that were covered by at least one short read. We considered a MAG detected in a metagenome if the detection value was > 0.25, which is an appropriate cutoff to eliminate false-positive signals in read recruitment results for its genome.

We annotated the MAGs of interest with DRAM, a genome annotation tool that provides metabolic profiles for each of the input MAGs. More detailed methods are provided in Shaffer, Borton et al. [45]. Briefly, Prodigal is used to detect open reading frames (ORFs) and predict their amino acid sequences, supporting all genetic codes as defined on NCBI. DRAM then searches all amino acid sequences against KEGG, UniRef90, and MEROPS using MMseqs2, with the best hits (defined by bit score with a default minimum threshold of 60) reported for each database [46]. We also used DRAM to perform HMM profile searches of the Pfam database and HMMER3 for dbCAN and VOGDB, with coverage length >35% of the model and e-value <10^-15^ to be considered a hit [40].

### Soil chemical analyses

To characterize the soil chemical environment of the samples, we measured soil pH, ferrous (Fe(II) and ferric (Fe(III) iron), and soil inorganic N concentrations as follows. Concentrations of Fe(II) and Fe(III) in 0.5 N HCl was measured as a representative anaerobic microsite in the samples. Samples were thawed and homogenized in an anaerobic chamber (Coy Labs, 2-5% H2 with N2 balance and Pd catalyst). A subsample (1 g) of the homogenized samples was placed into three 15 mL centrifuge tubes with 10 mL of 0.5 N HCl, removed from the anaerobic chamber, and shaken on an orbital shaker for 1 hour. The tubes were then centrifuged at 5000 rpm for 5 minutes, and Fe(II) was quantified with a UV‒Vis spectrophotometer (Thermo Genesys 10S) using the ferrozine method [47]. For quantification of Fe(III) concentrations, iron in the extract solutions was reduced to Fe(III) with hydroxylamine hydrochloride and measured with a spectrophotometer. The iron analysis results were normalized to the dry weight of the samples. Soil pH was measured in suspensions of fresh soil (2 g) and deionized water/0.01 M calcium chloride (10 mL) and were agitated for 20 minutes. Solutions were agitated on an orbital shaker, and pH was measured using a single junction general-purpose pH electrode connected to a PC-450 meter (Oakton). Soil inorganic N (NO_3_N and NH_4_N) was extracted with 1 M KCl (1:5 soil/solution ratio), filtered through a Whatman no. 42 paper filter, and analyzed with a continuous flow colorimetric analyzer (Lachat Instruments). Soil moisture was determined by weighing 10 g soil and drying at 105°C until constant weight. In-season soil inorganic N was expressed as a concentration in soil (μg kg^−1^).

### Statistical analyses

Differences between groups were analyzed by ANOVA unless stated otherwise. P values of less than 0.05 were considered statistically significant. We used RStudio v1.3.1093153 to visualize MAG detection patterns (https://www.rstudio.com/products/rstudio/) using pheatmap (pretty heatmaps) v1.0.12, ggplot2 v3.3.5 (https://ggplot2.tidyverse.org/), forcats v0.5.1 (https://forcats.tidyverse.org/), dplyr v1.0.8 (https://dplyr.tidyverse.org/), and ggpubr v0.4.0 (https://CRAN.R-project.org/package=ggpubr) [48]. We used multiple linear regression analysis to predict the MAG detection rate across land legacy plots for soil pH, NO_3_N, NH_4_N, Fe, and their interactions.

### Data availability

We uploaded our metagenome raw sequencing data to the SRA under NCBI BioProject PRJNA855256. All other analyzed data in the form of databases and fasta files are accessible at figshare 10.6084/m9.figshare.24243646.

## Results and discussion

### Genome-resolved analysis of metagenomes of field and monolith cores yields ammonia-oxidizing microorganism MAGs

We analyzed a total of 17 field cores (1 sample did not meet the minimum QC criteria for shotgun metagenomic sequencing) and 36 monolith samples (Supplementary Table S1). Of the 42,384,585 ± 21,156,189 metagenomic reads, an average of 39,720,343 ± 19,669,881 reads per sample passed the quality control criteria and were used for 4 coassemblies (Supplementary Table S2). After automatic and manual curation of the bins, we resolved 626 nonredundant MAGs (completion ≥70%, redundancy <10%), an average of 455 contigs with an N50 of 16,405 ± 21,253. The resolved MAGs were annotated to the domains Bacteria (n=565), Archaea (n=39), and Unknown (n=22). The genomic completion estimates for our MAGs averaged 82.1%, and the most abundant microbial populations were resolved to the phyla Actinobacteriota (n=243), Acidobacteriota (n=96), Proteobacteria (n=90), Verrucomicrobiota (n=33) and Methylomirabilota (n=31) (Supplementary Table S3).

### Prevalence of ammonia-oxidizing archaea and bacteria in drier conditions and differentiation of land legacy

Our genomic collection included 22 ammonia-oxidizing archaea (AOA) and 6 ammonia-oxidizing bacteria (AOB) that contained the complete set of genes for ammonia oxidation. To the best of our knowledge, these MAGs represent the first genomic evidence of putative AOA and AOB inhabiting the surface of dry soil conditions. MAGs resolved to AOA had an average genome size of 2.34 Mbp had a GC content ranging from 42.8% to 29.9%. The MAGs that were resolved to AOB had an average of 2.55 Mbp and CG content ranged from 59.3% to 56.5%. Five of the AOA MAGs were resolved to the genus *Nitrososphaera*, 7 AOA MAGs corresponded to lineages from the genus *TA-21*, and 1 AOA MAG was resolved to the genus *UBA10452*. The remaining AOA MAGs (n=9) from the Thermoproteota phylum corresponded to lineages within the family Nitrososphaeraceae. The phylogenetic assignment of all 6 AOBs was resolved to the genus *Nitrospira* (Figure 2, Table 1). We identified a total of 71,791 genes in the 22 AOA and 6 AOB MAGs (Supplementary Table S4). On average, the proportion of unknown function was 68% for the AOA MAGs and 63% for the AOB MAGs, reflecting our greater lack of functional understanding of the taxonomic group of ammonia-oxidizing microbial populations.

**Figure 2.**
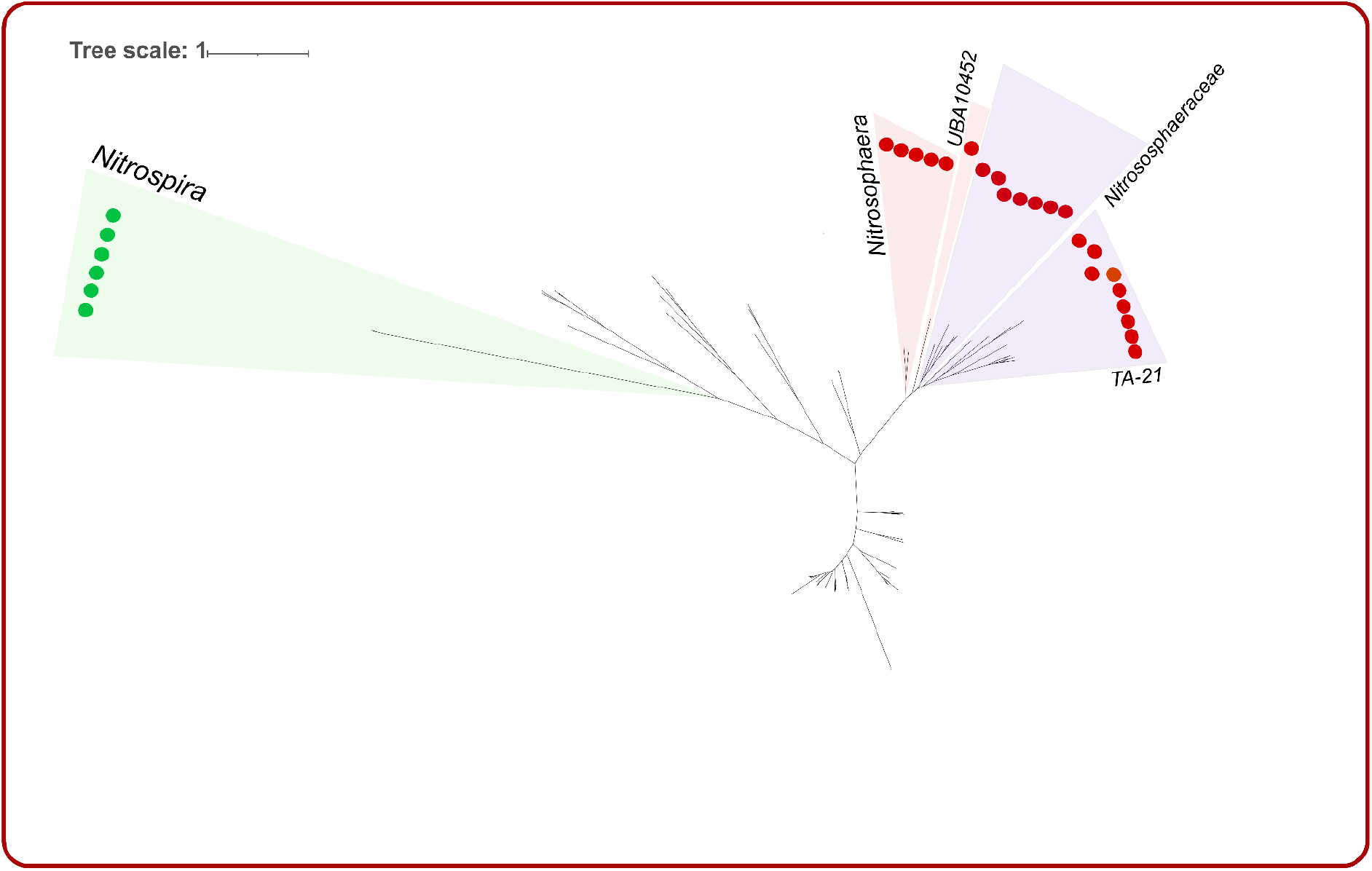
Phylogenetic tree of the 22 ammonia-oxidizing archaea (AOA) and 6 ammonia-oxidizing bacteria (AOB) MAGs. AOB MAGs were clustered together on the branch of *Nitrospira*, while AOA MAGs were assigned to *Nitrosophaera*, *UBA10452, Nitrososphaeraceae*, and *TA-21*.

**Table 1.** Assembly statistics of the 22 ammonia-oxidizing archaea (AOA) and 6 ammonia-oxidizing bacteria (AOB) MAGs, including total length, domain, family, and genus.

| MAGs | total_length | domain | family | genus |
| --- | --- | --- | --- | --- |
| Ar_MAG_001 | 2,216,557 | Archaea | Nitrososphaeraceae | Nitrososphaera |
| Ar_MAG_002 | 1,691,400 | Archaea | Nitrososphaeraceae | Nitrososphaera |
| Ar_MAG_003 | 2,985,253 | Archaea | Nitrososphaeraceae | TA-21 |
| Ar_MAG_004 | 2,261,092 | Archaea | Nitrososphaeraceae | - |
| Ar_MAG_005 | 2,185,676 | Archaea | Nitrososphaeraceae | - |
| Ar_MAG_006 | 4,203,210 | Archaea | Nitrososphaeraceae | TA-21 |
| Ar_MAG_007 | 2,258,495 | Archaea | Nitrososphaeraceae | - |
| Ar_MAG_008 | 2,492,023 | Archaea | Nitrososphaeraceae | - |
| Ar_MAG_009 | 1,988,562 | Archaea | Nitrososphaeraceae | Nitrososphaera |
| Ar_MAG_010 | 1,557,796 | Archaea | Nitrososphaeraceae | Nitrososphaera |
| Ar_MAG_011 | 2,700,998 | Archaea | Nitrososphaeraceae | Nitrososphaera |
| Ar_MAG_012 | 1,595,809 | Archaea | Nitrososphaeraceae | TA-21 |
| Ar_MAG_013 | 2,003,530 | Archaea | Nitrososphaeraceae | TA-21 |
| Ar_MAG_014 | 2,041,095 | Archaea | Nitrososphaeraceae | TA-21 |
| Ar_MAG_015 | 1,919,785 | Archaea | Nitrososphaeraceae | TA-21 |
| Ar_MAG_016 | 1,959,807 | Archaea | Nitrososphaeraceae | TA-21 |
| Ar_MAG_017 | 1,699,605 | Archaea | Nitrososphaeraceae | UBA10452 |
| Ar_MAG_018 | 3,702,399 | Archaea | Nitrososphaeraceae | - |
| Ar_MAG_019 | 3,161,044 | Archaea | Nitrososphaeraceae | - |
| Ar_MAG_020 | 3,657,672 | Archaea | Nitrososphaeraceae | - |
| Ar_MAG_021 | 2,385,568 | Archaea | Nitrososphaeraceae | - |
| Ar_MAG_022 | 728,568 | Archaea | Nitrososphaeraceae | - |
| Ba_MAG_023 | 1,726,077 | Bacteria | Nitrospiraceae | Nitrospira |
| Ba_MAG_024 | 3,180,767 | Bacteria | Nitrospiraceae | Nitrospira |
| Ba_MAG_025 | 2,620,829 | Bacteria | Nitrospiraceae | Nitrospira |
| Ba_MAG_026 | 3,050,531 | Bacteria | Nitrospiraceae | Nitrospira |
| Ba_MAG_027 | 2,102,900 | Bacteria | Nitrospiraceae | Nitrospira |
| Ba_MAG_028 | 2,640,344 | Bacteria | Nitrospiraceae | Nitrospira |

In this study, assessing the abundance of AOO MAGs in different environments provides an opportunity to investigate the association between AOA and AOB with precipitation and land legacy (Figure 3). We showed that there was a statistically significant difference in the abundance of AOA and AOB MAGs between the eastern and western Kansas soils (PERMANOVA: Pseudo-F= 9.0280, p= 0.001, Figure 3). We also showed a statistical significance between the different land use legacies (PERMANOVA: Pseudo-F= 4.4636, p= 0.001, Figure 3) as well as an interactive effect between location and land use legacy (PERMANOVA: Pseudo-F= 4.2306, p= 0.001, Figure 3). We observed that AOA and AOB MAGs were distributed in 3 distinct clusters based on the interaction of the environmental variables. Cluster-Ag was highly detected in the agricultural soils and tapered off in detection through the native and postagricultural soils. MAGs from Cluster-Ag were also sparingly detected in field cores from WKS. MAGs from Cluster-Low-Detection were the least detected in the samples among all the MAGs, with some of the MAGs detected in the post-Ag soils. We observed that the last cluster (Cluster-Native) had a reverse detection pattern in the MAGs compared to that of Cluster-Ag. There was a total of 9 MAGs in this cluster (Figure 3).

**Figure 3.**
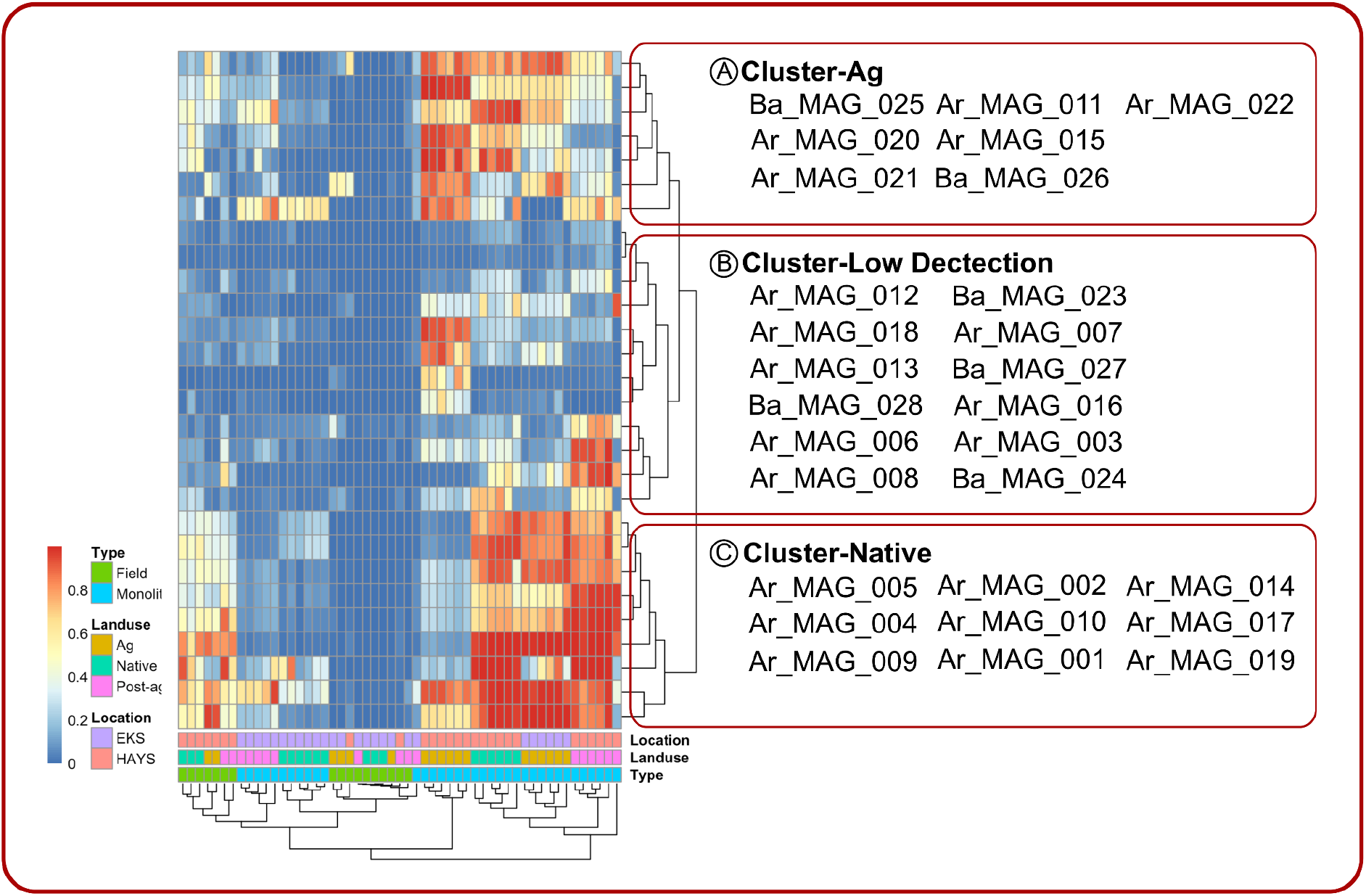
MAGs are clustered into three distinct clusters: Cluster-Ag, Cluster-Low Detection, and Cluster-Native. The clustering of the MAGs was mainly driven by the detection of the AOA and AOB MAGs in the drier locations (HAYS).

MAGs were detected more frequently in the monolith than in the field cores, which suggests enrichment of ammonia-oxidizing microorganisms, providing an opportunity to assemble the genomes of these AOOs. We also postulated that the drying of the soil in the monoliths provided an environment more conducive for the AOOs to thrive [49,50]. Among the field core samples, we also observed that AOO MAGs were highly detected in WKS but almost undetectable in EKS, consistent with the hypothesis that AOA and AOB in our study were detected in drier conditions. Consistent with our findings, there are reports that soil moisture showed a negative correlation with the abundance of AOA in tall grass prairies [51]. In previous reports, archaeal amoA genes were found to be more abundant in drier soil, but within these microenvironments, they were 40% more abundant in water-filled space, which suggested that reduced oxygen levels decreased the growth of AOA [52]. Taken together, our results suggest that AOA and AOB MAGs are more prevalent in drier soil conditions [53]. We further performed correlation analyses to identify the relationships between the detection levels of AOO MAGs and various soil variables (soil pH, iron(III) oxide, nitrate, and ammonia). As expected, we observed a decrease in the detection levels of AOO MAGs with the increase in iron(III) oxide in the soil (R = –0.28, P < 0.001) (Figure 4A, Supplementary Figure S1). Similarly, we showed that the detection of AOO MAGs was lower with higher soil pH (R = –0.26, P < 0.001). We also observed a positive correlation between AOO MAG detection levels and nitrate (R = 0.37, P < 0.001) and a marginal positive correlation with ammonia levels (R = 0.082, P = 0.075) in the soil (Figure 4A, Supplementary Figure S1).

**Figure 4.**
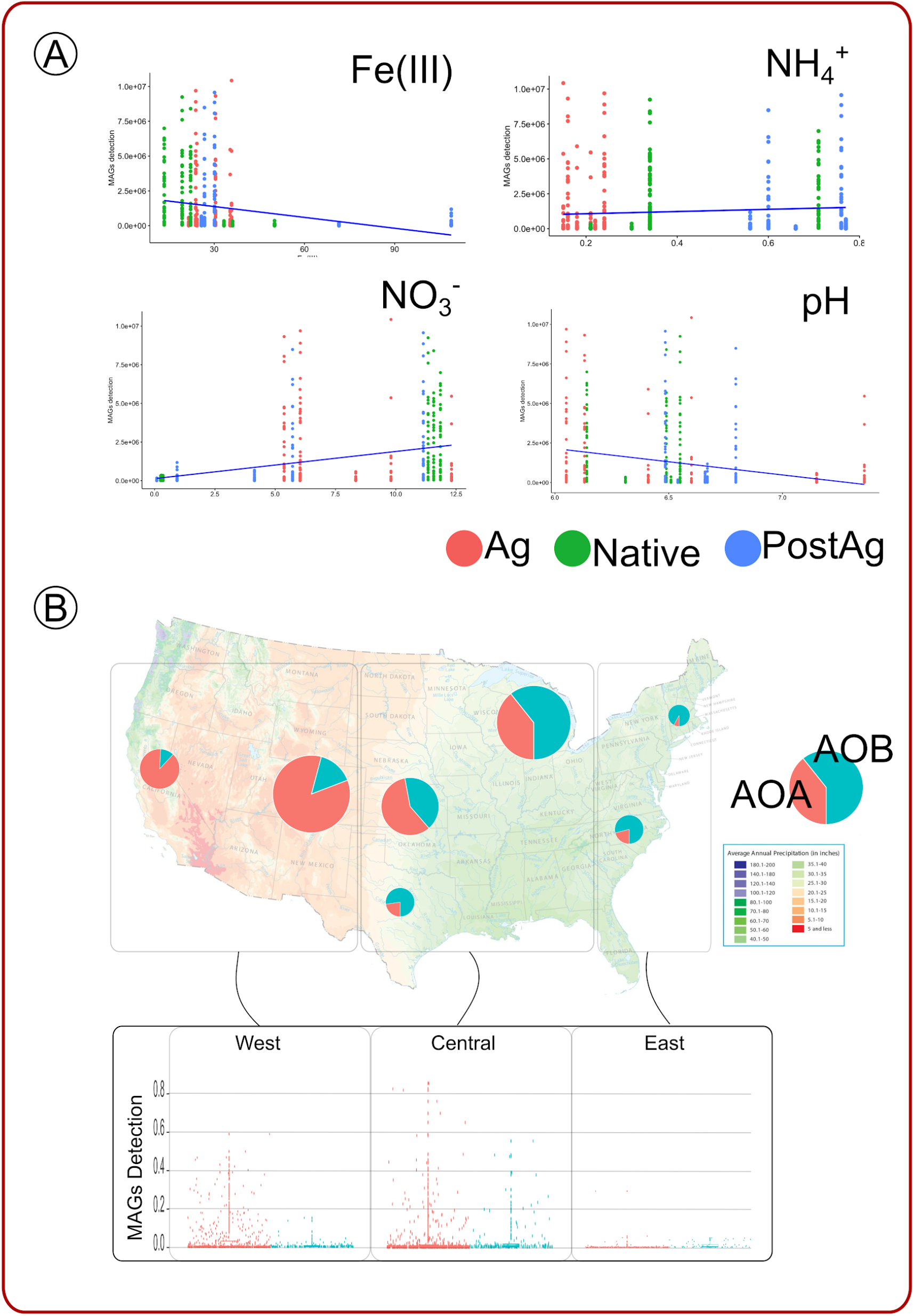
(A) The correlation between the detection levels of AOO MAGs and various soil variables (soil pH, iron(III) oxide, nitrate, and ammonia). The detection of AOO MAGs was negatively correlated with an increase in iron(III) oxide in the soil and higher soil pH. There were positive correlations between AOO MAG detection levels and nitrate and a marginal positive correlation with ammonia levels in the soil. (B) The distribution of the 28 nonredundant AOO MAGs with precipitation levels across the country with 3 US different regions –West, Central, and East. AOA and AOB MAGs were mostly detected in the central and western regions but were barely detected along the east coast.

To further understand the distribution of AOO MAGs across the precipitation gradient in the United States, we mapped 70 downloaded metagenomic datasets from soil profiles (Supplementary Table S5) to the AOA and AOB MAGs. The 28 nonredundant AOO MAGs had a distinct distribution across the different landscapes of the United States, which correlated with precipitation levels across the country (Figure 4B). We categorized the locations where the metagenomes were sourced into 3 US regions –West, Central, and East. We noticed that AOA and AOB MAGs were mostly detected in the Central (AOA: 50.1%, AOB: 85.4%) and Western regions (AOA: 49.4%, AOB: 12%) but were barely detected along the East coast (AOA: 0.35%, AOB: 1.85%) (Figure 4B). We also downloaded the US precipitation map (https://gisgeography.com/us-precipitation-map/) and showed that the AOOs were mostly detected in the drier Central area where they are dominated by AOB MAGs (Figure 4B).

### Stress response functional potential is critical for ammonia-oxidizing archaea and bacterial resilience

Our metagenomic data showed that the resolved AOA and AOB MAGs harbored a total of 244 and 71 gene functions based on Cluster of Orthologous (COG) functions that were involved in ammonia-oxidation (Supplementary Table S6). We detected several genes that were related to (1) nitrite reductase (NADH) small subunit, (2) hydroxylamine dehydrogenase, and (3) methane/ammonia monooxygenase subunit A, supporting our conclusion that our resolved AOA and AOB MAGs possessed key enzymes essential to ammonia oxidation and have the potential to be involved in microbial oxidation of ammonia to nitrite [54]. Nitrite reductase is a key enzyme involved in denitrification and plays a critical role in the N cycle [55], while hydroxylamine dehydrogenase is another important enzyme identified in bacteria involved in aerobic ammonia oxidation [56,57]. Methane/ammonia monooxygenase subunit A belongs to proteins well identified in methanotrophs and ammonia oxidizers [58] and catalyzes the first and rate-limiting step in the nitrification process [8].

We further observed a prevalence of carbon-fixing genes in AOO MAGs (Supplementary Table S6). AOA MAGs (68.03%) showed a higher number of carbon-fixing genes as compared to AOB MAGs (31.97%). Furthermore, most of the AOA MAGs showed the potential to produce phenols, which might be due to the antioxidant activity in the AOA and suggested to be generated by oxidative stress [59]. We deduced that most of the AOA and AOB could use the 3-hydroxypropionate bicycle pathway to fix carbon [60]. The 3-hydroxypropionate is a novel pathway for autotrophic carbon fixation and was first identified in *Chloroflexus aurantiacus* [61]. Later, it was identified in different members of the domain Archaea [61]. Our study emphasizes the capability and potential mechanism of AOA and AOB to fix carbon.

The availability of O_2_ is generally less in wet soil than in dry soil [62]. Therefore, it is interesting to note that the AOA and AOB are facilitating higher fixation potentials in dry soil. The AOA and AOB contributing to the N cycle could also be important to the global carbon cycle. AOA from are canonically autotrophs, with some evidence that they could also be mixotrophic [63–65]. However, we are still unclear the extent of the functional potential in AOA and AOB [60]. Our study potentially confirms that the AOA and/or AOB have a substantial contribution to ammonia oxidation and carbon fixation in terrestrial environments.

Similarly, we identified 303 and 89 stress response genes based on COG functions in the AOA and AOB MAGs, respectively (Supplementary Table S6). Some of the gene functions that we observed were (1) Desiccation stress tolerance protein, LEA/WHy domain (LEA), (2) predicted membrane GTPase TypA/BipA involved in stress response (TypA), and (3) nucleotide-binding universal stress protein, UspA family (UspA). Desiccation stress tolerance proteins with the LEA/WHy domain are a part of defense mechanisms in bacteria under abiotic stresses [66]. The predicted membrane GTPase TypA/BipA involved in the stress response (TypA) [67] and nucleotide-binding universal stress protein UspA family (UspA) [68] impart resistance to bacteria. The GTPase TypA/BipA involved in the stress response (TypA) is reported to have chaperone-like activity [67]. The nucleotide-binding universal stress protein UspA family (UspA) is critical for bacterium survival during cellular growth arrest [68]. Our analysis of stress response genes from AOO MAGs indicated that stress responses were likely key functional potentials contributing to archaeal and bacterial prevalence under drier conditions.

### Differential strategies of ammonia-oxidizing archaea and bacteria

To assess whether our AOA and AOB MAGs adopted different survival strategies, we used Distilled and Refined Annotation of Metabolism (DRAM) to profile and compare the metabolic potentials harbored in the AOO MAGs. We noticed that ammonia-oxidizing genes were highly detected in the AOA MAGs, while the AOB populations showed high detection of aerobic-specific ammonia-oxidizing genes. Furthermore, we also noticed that AOA MAGs had the functional potential to produce nitric oxide from nitrite metabolism (Supplementary Table S7). We further examined the sulfur metabolism potential of the MAGs and noticed that only AOB populations had the potential to oxidize thiosulfate to sulfate using the SOX complex, suggesting that the AOB MAGs might have an alternative energy pathway to ammonia oxidation (Supplementary Table S). *Nitrospira* can oxidize sulfur [69]. The gene set involved in sulfur oxidation consists of sat genes encoding a sulfate adenylyl transferase, aprAB genes encoding an adenylyl sulfate reductase, qmoABC genes encoding a complex involved in electron transfer, and an array of dsr genes whose encoded proteins catalyze reverse reactions of sulfate reduction [70]. These sulfur-oxidizing bacteria (SOB) typically have dsrEFH genes, whose products react with DsrC in the process of sulfur oxidation [71].

We used the AOA and AOB MAGs to predict the mechanism of AOO MAGs in our study. We observed that all AOO MAGs contained modules for energy metabolism (citrate cycle, dicarboxylate-hydroxybutyrate cycle, Embden–Meyerhof–Parnas (EMP) pathway, glyoxylate cycle, and hydroxypropionate-hydroxybutyrate cycle). We also noticed that the AOA and AOB populations in our study might harbor alternative strategies for energy transformation. AOB MAGs contained genes that had metabolic potential to transform glucose-6P to glyceraldehyde-3P and pyruvate for their energy requirement using the Entner–Doudoroff (ED) pathway, while the AOA MAGs completely lacked the genes for this metabolic potential (Figure 5). Flamholz et al. argued that the choice between the EMP and ED pathways is a tradeoff between ATP yield and protein costs [72]. In grassland prairie where there are frequent dry and wet conditions, the utilization of the EMP pathways is particularly advantageous for microorganisms with a facultatively anaerobic lifestyle that gain energy mainly by fermentative metabolism [72]. We further observed that the majority of the AOA MAGs (14 out of 22 MAGs) harbored functional pathways for methanogenesis, whereas this potential function was completely missing in all AOB MAGs. Methanogenic AOA can produce methane (CH_4_) and have the potential to contribute significantly to global climate change [73]. Although the capacity for mixotrophy has been proposed for soil AOA [16], the mechanism is not well understood. Methanogenesis is only energetically favorable for AOA under highly reduced conditions; i.e., waterlogged for long enough to deplete all other electron acceptors to generate a proton gradient to run ATP synthase. While the conditions needed for AOA methanogenesis are the opposite for conditions for ammonia oxidation, here, we showed genomic evidence for the first time of the potential mixotrophic ability of AOA for methanogenesis as well as ammonia oxidation, fueling the competitiveness of AOA over AOB under specific soil conditions.

**Figure 5.**
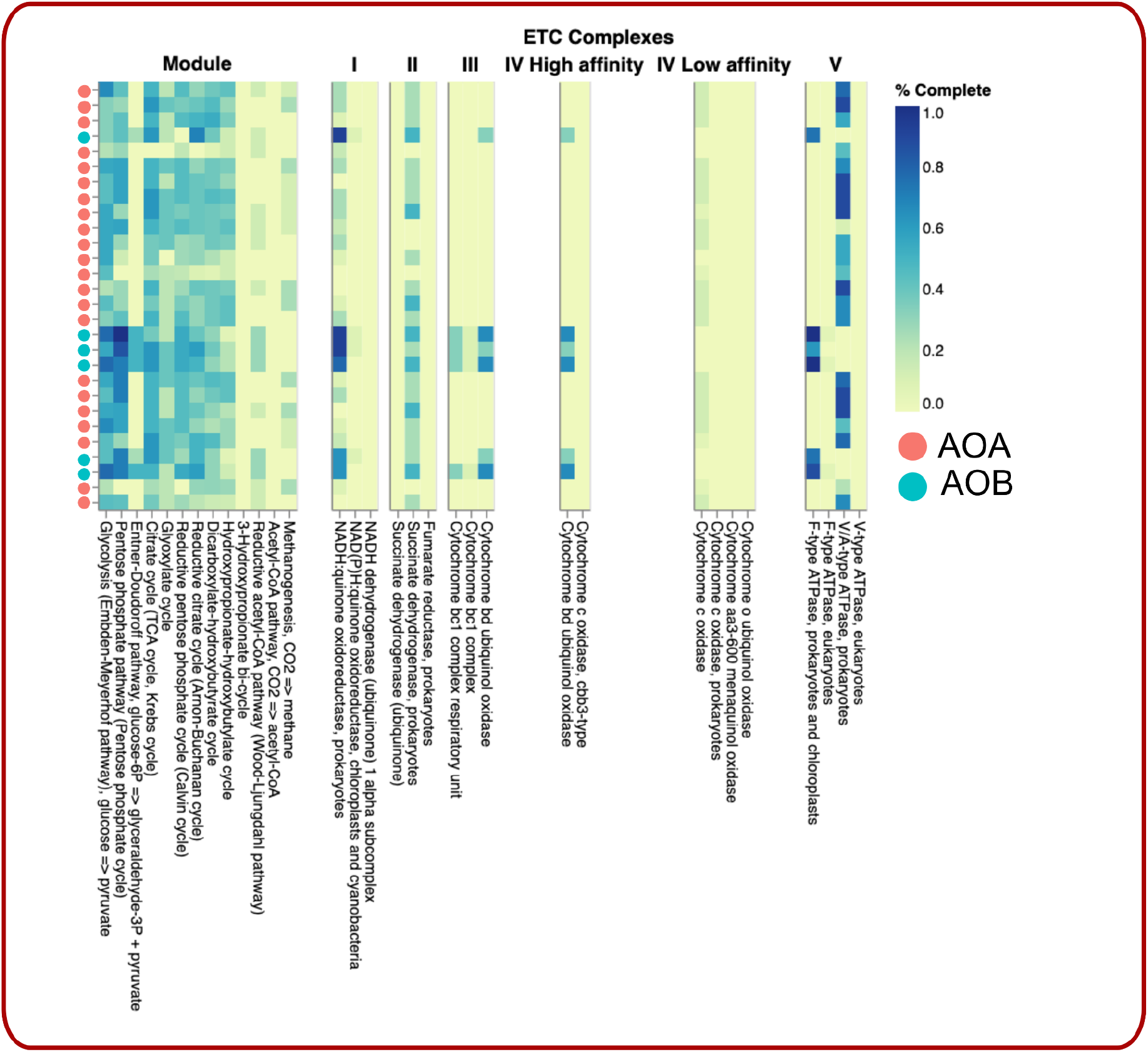
Distilled and Refined Annotation of Metabolism (DRAM) analysis compared the metabolic potentials harbored in the AOB and AOA MAGs. AOB MAGs contained genes of the Entner–Doudoroff (ED) pathway, while the AOA MAGs completely lacked the genes for this metabolic potential. All AOA MAGs were missing canonical membrane-bound electron transport chains, and most AOB MAGs had a complete complex III module.

All AOA MAGs were missing canonical membrane-bound electron transport chains, as complexes II, III and IV were either absent or incomplete (Figure 5). We also noticed that most AOB MAGs had a complete complex III module (cytochrome bc1 complex, cytochrome bc1 complex respiratory unit, cytochrome bd ubiquinol oxidase). The F-type ATPase was present in all AOB MAGs but missing in the AOA populations. On the other hand, only AOA MAGs harbored A/A-type ATPase. The key enzymes in energy conservation are the archaeal A_1_A_O_ ATP synthases, a class of ATP synthases distinct from the F_1_F_O_ ATP synthase ATP synthase found in bacteria, mitochondria, and chloroplasts, and the V_1_V_O_ ATPases of eukaryotes [74]. F-type ATPases are located in the membranes of bacteria, chloroplasts, and mitochondria and catalyze the hydrolysis or synthesis of ATP coupled with H+ (or Na+) transport across a membrane [75]; thus, it is accepted that aerobic organisms use the enzyme mainly for synthesizing ATP [75]. The archaeal A/A-type ATPases have nine subunits, whereas the bacterial F-ATPases have eight different subunits [76]. Structurally, A_1_A_0_ ATP synthases are unique from F_1_F_O,_ as they have an optimized P-loop design in the catalytic A subunit, which allows the enzyme to catalyze ATP along with UTP and GTP [74]. Additionally, A/A-type ATPases possess additional α-helices at the C-terminus of subunit A, which is absent in the F-ATPase [77]. The major strategy of archaeal A/A-type ATPases has been to tighten up the coupling mechanism and prevent any leakage of coupling ions through the ATP synthase motor [76]. The A_1_A_0_ ATP synthases also rely on sodium and/or proton gradients to encounter high temperatures and low-energy environments [76]. The assessment of the metabolic potentials of bacteria and archaea suggested different survival strategies adopted by AOA and AOB [78].

## Conclusions

Our study provides new knowledge of how ammonia-oxidizing archaea and ammonia-oxidizing bacteria live in soil, contributing to important N-transforming processes, and are impacted by varying environmental factors such as precipitation and land legacies. We identified MAGs that represented the first genomic evidence of AOA and AOB in dry soil conditions. We showed that AOA and AOB had a higher presence in drier conditions, and land legacies could regulate the differential abundance of the AOA and AOB. We believe it is even more important to dissect the functional potentials of the AOA and AOB to gauge more insights into the contribution of the AOA and AOB populations to the N-transforming processes under variable soil environmental conditions. We have demonstrated that the AOA and AOB use stress response genes, differential metabolic functional potentials, and subtle population dynamics under different environmental parameters. The knowledge from this study will help us to understand how future climatic changes such as drought-like conditions individually or in the combination of native, agricultural, and postagricultural land legacies could have an impact on AOA and AOB driving N transformations in soil.

## List of abbreviations

N_2_O: Nitrous Oxide; AOA: Ammonia-Oxidizing Archaea; AOB: Ammonia-Oxidizing Bacteria; MAGs: Metagenome Assembled Genomes; ATPase: Adenosine Triphosphatase; N: Nitrogen; GHGs: Greenhouse gases; AOOs: Ammonia-Oxidizing Microorganisms; WKS: Western Kansas; EKS: Eastern Kansas; ORFs: Open Reading Frames; COGs: Clusters of Orthologous Groups; ANI: Average Nucleotide Identity; DRAM: Distilled and Refined Annotation of Metabolism; SOB: Sulfur-Oxidizing Bacteria; EMP: Embden–Meyerhof–Parnas; ED: Entner– Doudoroff.

## Declarations

### Ethics approval and consent to participate

Samples were collected on experimental grounds belonging to Kansas State University and University of Kansas. No permission was necessary to collect the samples used in this study.

### Consent for publication

Not Applicable.

### Availability of data and material

We uploaded our metagenome raw sequencing data to the SRA under NCBI BioProject PRJNA855256. All other analyzed data in the form of databases and fasta files are accessible at figshare 10.6084/m9.figshare.24243646.

### Authors’ contributions

S.S. and A.K. conceptualized, conducted the experiments, performed the data analysis, and wrote the manuscript. P.M.H. conducted the experiments, analyzed the data, and wrote the manuscript. K.W., C.H., and N.R. conducted the experiments and analyzed the data. Q.R. analyzed the data. W.K., and L.F.T.deS. conducted the experiments. T.D.L., M.V.M.S., C.W.R, L.H.Z, B.A.S. conducted the experiments, and wrote the manuscript. S.T.M.L. conceptualized, performed the data analysis, supervised, was responsible for resource acquisitions, and wrote the manuscript. All the authors contributed to the article and approved the submitted version.

## Acknowledgments

We appreciate the Kansas Medical Center Genomics Core for the metagenomic data generated.

## Supplementary Figures and Tables

Supplementary Figure S1. Correlation analyses between the detection levels of AOO MAGs and various soil variables (soil pH, iron(III) oxide, nitrate, and ammonia).

Supplementary Table S1. A list of 17 field cores and 36 monolith samples was included in the study with the types, locations, and land use.

Supplementary Table S2. Assemble statistics of the samples.

Supplementary Table S3. Nonredundant MAGs from field cores and monolith samples.

Supplementary Table S4. Gene calls from 22 AOA and 6 AOB MAGs.

Supplementary Table S5. Seventy metagenomic datasets were downloaded to map soil profiles to the AOA and AOB MAGs.

Supplementary Table S6. Gene functions of AOA and AOB MAGs that are involved in ammonia oxidation, carbon fixation, and stress responses along with the detection numbers.

Supplementary Table S7. Features from DRAM analysis include carbon utilization, transporters, energy, organic nitrogen, carbon utilization, rRNA, and tRNA.

