## Supplementary figures and images for "Ammonia-oxidizing archaea and bacteria differentially contribute to ammonia oxidation in soil under precipitation gradients and land legacy"

### Supplementary Figure1

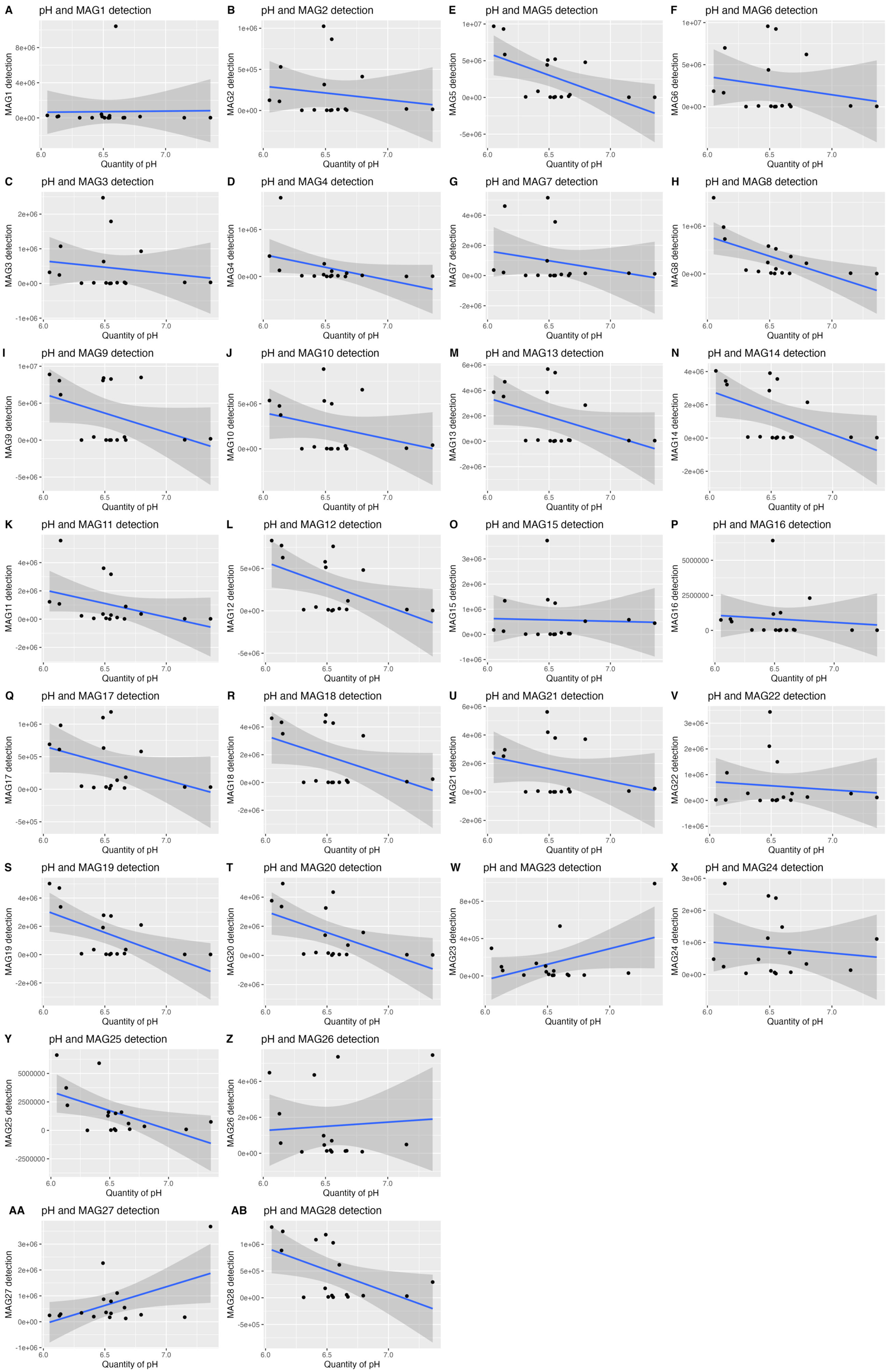

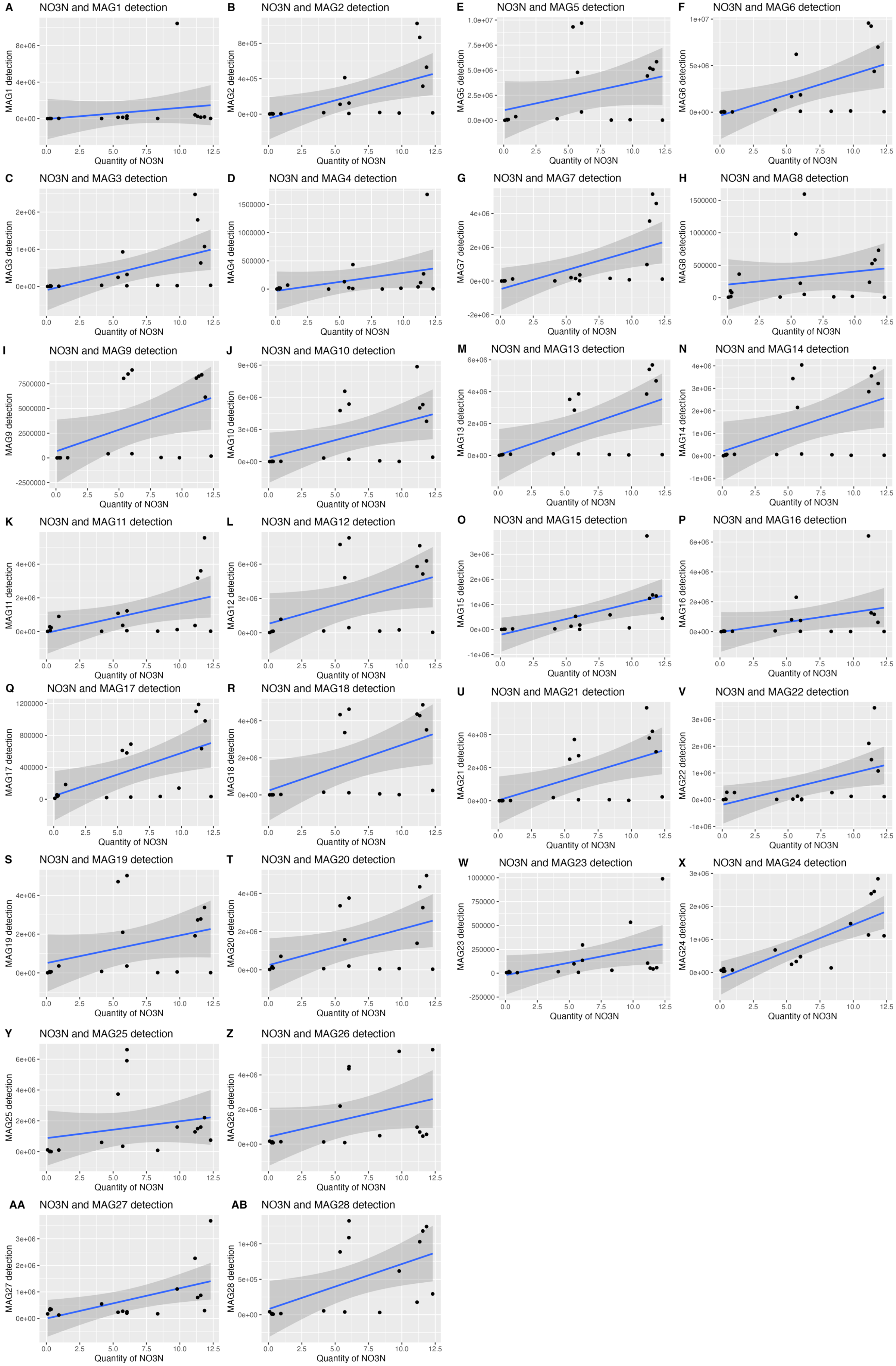

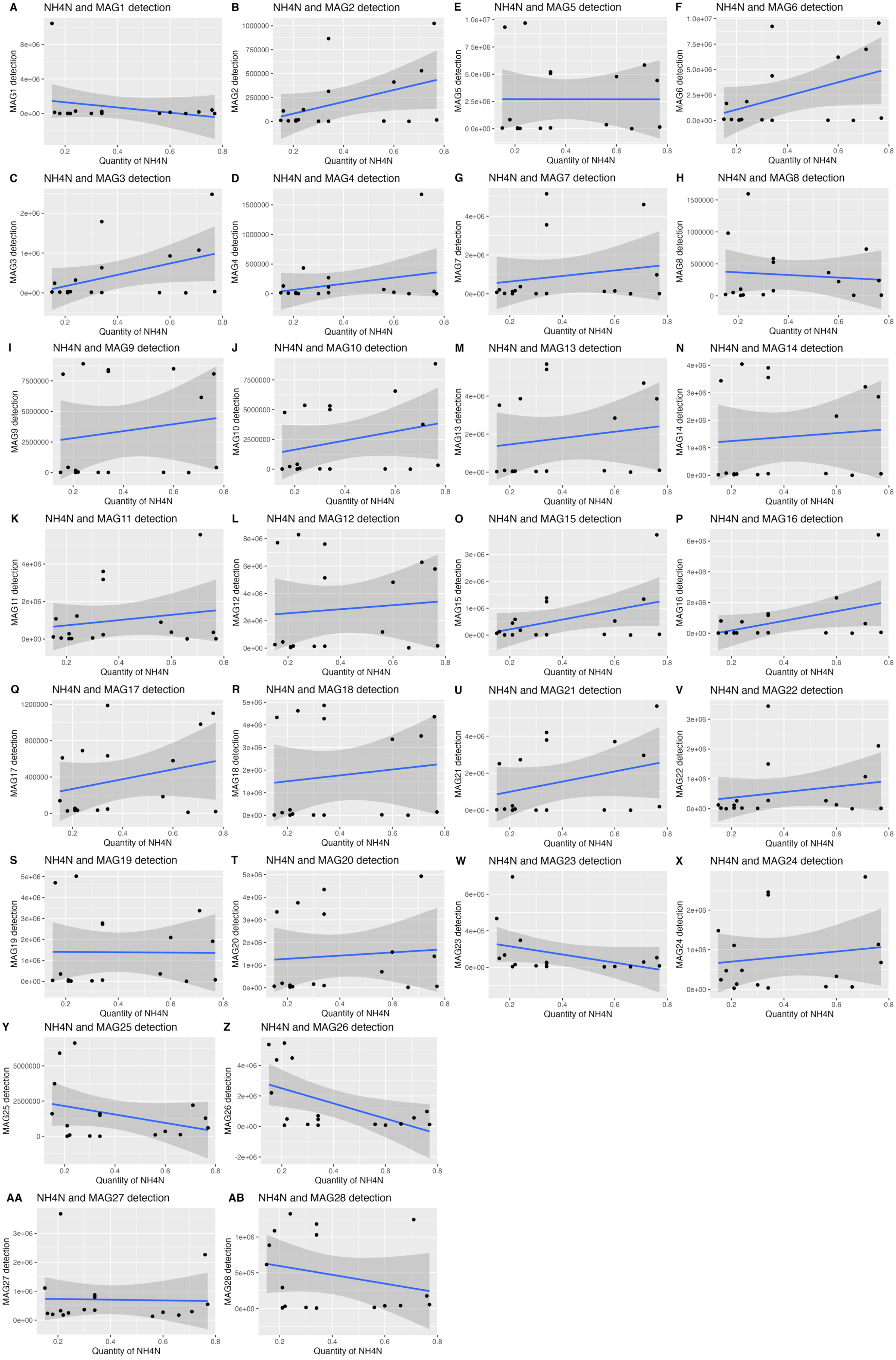

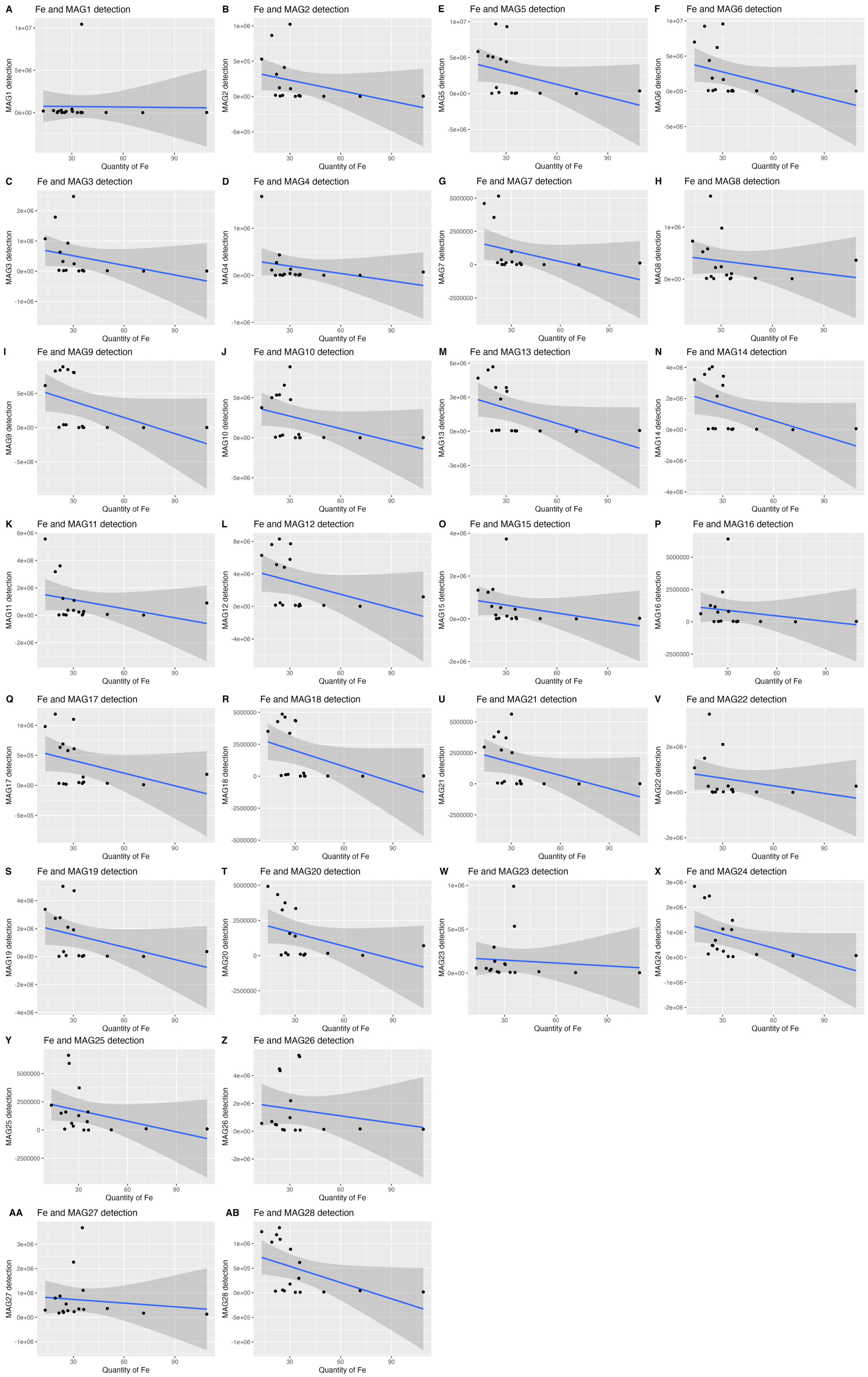
